# Germline burden of rare damaging variants negatively affects human healthspan and lifespan

**DOI:** 10.1101/802082

**Authors:** Anastasia V. Shindyapina, Aleksandr A. Zenin, Andrei E. Tarkhov, Peter O. Fedichev, Vadim N. Gladyshev

**Author notes:** Correspondence and requests for materials should be addressed to V.N.G. and P.O.F. These authors also contributed equally to this work (Anastasia V. Shindyapina, Aleksandr A. Zenin). These authors contributed equally to this work (Peter O. Fedichev, Vadim N. Gladyshev).

## Abstract

Genome-wide association studies often explore links between particular genes and phenotypes of interest. Known genetic variants, however, are responsible for only a small fraction of human lifespan variation evident from genetic twin studies. To account for the missing longevity variance, we hypothesized that the cumulative effect of deleterious variants may affect human longevity. Here, we report that the burden of rarest protein-truncating variants (PTVs) negatively impacts both human healthspan and lifespan in two large independent cohorts. Longer-living subjects have both fewer rarest PTVs and less damaging PTVs. In contrast, we show that the burden of frequent PTVs and rare non-PTVs is less deleterious, lacking association with longevity. The combined effect of rare PTVs is similar to that of known variants associated with longer lifespan and accounts for 1 − 2 years of lifespan variability. We further find that somatic accumulation of PTVs accounts for a minute fraction of mortality and morbidity acceleration and hence provides little support for its causal role in aging. Thus, damaging mutations, germline and somatic, can only contribute to aging as a result of higher-order effects including interactions of multiple forms of damage.

## Introduction

Genome-wide association studies (GWAS) of human lifespan, including studies examining extreme longevity, parental survival, and healthspan, produced a number of gene variants associated with human longevity. For example, GWAS on centenarians consistently demonstrate the loci near *LPA* and *APOE, FOXO3A, HLA-DQA1* and *SH2B3* genes to be associated with longevity^1^. However, even in developed countries, centenarians currently represent less than 0.1% of the original cohort, and the genetic determinants responsible for the survival of general population remain poorly understood. Release of massive genotype and phenotype data by UK Biobank (UKB)^2^ inspired investigation of the relationship between genetics and longevity proxies, such as parental lifespan^3^ and healthspan within general population^4^. They confirmed most of the variants from centenarian studies and identified additional variants. However, the combined contribution of these variants could explain only a small part of lifespan heritability, at least as asserted from twin studies^5^. We hypothesized that the rest could be explained by the combined burden of rare damaging gene variants. Until very recently, only common variants could be probed in genetic studies due to sample size limitations. However, the large datasets such as gnomAD and UKB now allow assessing the effects of variants with minor allele frequency (MAF) lower than 0.1%.

These ultra-rare variants, most notably protein-truncating variants (PTVs), are known to be enriched for damaging alleles. They tend to have larger effect sizes and dramatically change gene expression and function. They are usually eliminated by purifying selection, but the small effective population size of the human population means that they are present in all human genomes. An inverse relationship between MAF and effect size was recently demonstrated for type II diabetes, an archetypal age-related disease^6^. Multi-tissue gene expression outliers were enriched with rare variants in the GTEx dataset^7^. Notably, PTVs represent a significant part of those variants. Most of the underexpressed outliers harbor rare variants and accordingly are subject to nonsense-mediated decay (NMD). Additionally, ExAC consortium demonstrated the nonsense variants with a high Combined Annotation Dependent Depletion score^8^, a widely used predictor of deleteriousness of single nucleotide variants, were enriched in singletons^9^. Although missense and non-coding variants may also be damaging, PTVs are substantially enriched for deleterious alleles. They also alter gene expression more dramatically than missense and untranslated region (UTR) variants^10^.

Increased rare PTV burden was shown previously to be associated with complex diseases, such as schizophrenia, epilepsy and autism^11–13^, whereas individual contributing genetic variants exhibited small effects. The burden of rare PTVs in genes intolerant to such variants (PI-PTVs) was tested for association across ExAC traits. For example, one study revealed a negative association with years of schooling (academic attainment) and a positive association with intellectual disability, autism, schizophrenia and bipolar disorder^14^. Notably, the age at enrollment was also negatively correlated with the burden of PI-PTVs referring to a possible association with lifespan. Overall, recent studies suggest that rare variants have a profound effect on complex traits and fitness. In this study, we focused specifically on the effect of germline PTV burden on longevity and disease-free survival, and estimated the effect of somatic PTV accumulation with aging on mortality and morbidity acceleration.

## Results

### Study design and data

We characterized the effect of ultra-rare damaging mutation burden on human traits associated with longevity. For the UK Brain Bank Network (UKBBN) postmortem samples, we run a survival analysis against the age at death^15^. For UKB subjects, we tested the effect of mutations on the remaining lifespan (or simply lifespan), that is survival within the available follow-up of eleven years. We also tested the effects of damaging gene variants on healthspan, that is the disease-free period (defined as the age when at least one of the following conditions is diagnosed for the first time: cancer, diabetes, myocardial infarction, congestive heart failure, chronic obstructive pulmonary disease, stroke, dementia, and death^4^). In addition, following the approach of^16^, we studied the effect of mutation load on parental survival (separately for the age at death for mothers and fathers), a useful longevity proxy in genetic studies.

We selected a cohort of 40, 368 individuals from UKB with sequenced exomes who selfreported ‘White British’ and were of close genetic ancestry based on a principal component analysis of their genotypes^2^. Of those, 21, 742 (54%) were males with mean age 58.1 years (*SD* = 7.9, age range 40.2 − 70.6) and 18, 626 (46%) were females with mean age 57 years (*SD* = 7.8, age range 40.1 − 70.4) at the time of assessment. In this cohort, 1, 122 subjects died during the follow-up period of 11 years (2005 − 2016), mostly of cancer (Table 1 and Table S1). The UKBBN cohort included 1, 105 subjects of European origin excluding cases of suicides, accidents and cases of death with no abnormalities detected. Of those, 489 (44%) were females with mean age of 71.2 years (*SD* = 18, 16 − 103 years) and 616 (56%) were males with mean age of 67.7 years (*SD* = 17, age range 17 − 105 years). The cause of death was reported for 359 individuals. Most of the participants in this study were diagnosed with neurodegenerative diseases, e.g. Alzheimer’s, Parkinson’s, and Pick’s diseases.

**Table 1:** Causes of death reported for 1, 122 and 359 subjects in UKB and UKBBN cohorts, respectively.

|  | UKB | UKBBN |
| --- | --- | --- |
| Neoplasm | 638(56.9%) | 20(5.6%) |
| Circulatory system | 208(18.5%) | 90(25.1%) |
| Respiratory system | 82(7.3%) | 171(47.6%) |
| Digestive | 47(4.2%) | 7(1.9%) |
| Nervous system | 43(3.8%) | 51(14.2%) |
| External | 35(3.1%) | — |
| Other (infections, congenital, endocrine, mental) | 69(6.1%) | 19(5.3%) |

As in^14^, we identified PTVs as variants that alter open reading frames of canonical transcripts, including splice donor/acceptors, stop codon gains and frameshifts. To address the relation between the frequency of mutations and their effects on lifespan, we binned gene variants from whole-exome sequencing (WES) datasets according to their frequency: 1) *MAF* < 10^−4^; 2) 10^−4^ < *MAF* < 10^−3^; 3) 10^−3^ < *MAF* < 0.01; 4) 0.01 < *MAF* < 0.2. We defined PTV exome burden as the total number of PTVs within the MAF bins (Figure S1 for PTV burden distribution in the MAF bins).

### Survival analysis

We examined the association of PTV load in the defined MAF bins against the selected longevity traits (i.e. survival in UKB and UKBNN, and healthspan and mother’s and father’s age at death in UKB) using Cox proportional hazards (PH) models. We used sex and genetic principal components as covariates to account for the effects related to the population heterogeneity. We additionally used age as a covariate for the UKB lifespan during the follow-up period.

The mortality and morbidity risk models returned the Cox regression parameters that were consistent with well-established mortality patterns. For example, the UKB survival model produced the regression coefficient Γ = 0.093 per year for the age of first assessment, very close to the mortality and morbidity acceleration rate (of approximately 0.09 per year in UKB cohort^4^) and hence consistent with the Gompertz mortality law. The Cox regression coefficient for sex was 0.47 for males in UKB and 0.29 in UKBBN. Under the constant mortality acceleration, this would correspond to approximately 3 − 5 years difference in life expectancy. Women in the UK (the population relevant to this study) live longer than males, although the gap between the sexes has decreased over time and is now 3.7 years^17^.

We found that, in both datasets, a higher burden of ultra-rare (*MAF* < 0.0001) PTVs negatively and significantly correlated with healthspan and lifespan (Figure 1). The proportional hazards effect estimations (sign and order of magnitude) were consistent, *β* = 0.046 and 0.014 per mutation, for lifespan and healthspan in UKB, respectively. Moreover, the Cox regression coefficients were very similar for UKBBN and UKB datasets, indicating consistency of the effect across populations despite differences in population structure and morbidity statistics (Table 1), tissue source, sequencing methods and variant calling pipelines.

**Figure 1:**
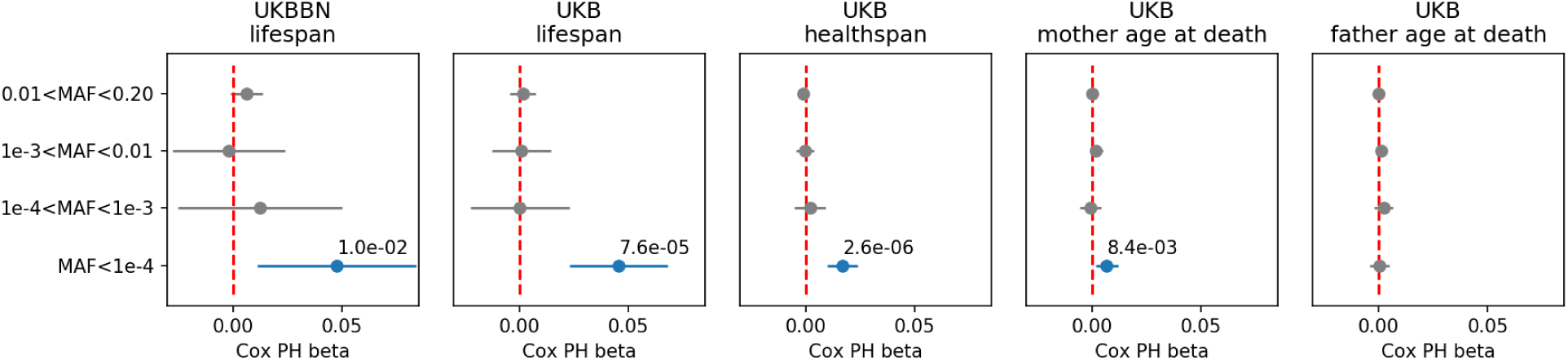
Burden of ultra-rare (*MAF* < 10^−4^) PTV variants is negatively associated with lifespan, healthspan and mother’s (but not father’s) age at death in UKBBN and UKB cohorts

We also observed a smaller but still significant effect of the ultra-rare PTV burden on mothers’ but not on fathers’ longevity in UKB. The effects on mother’s age at death was less significant than that on the survival and lifespan.

On average, we identified 6 (*SD* = 2.6) ultra-rare PTVs (*MAF* < 0.0001) per genome (Figure S1). To visualize the effects of such PTVs on survival, we split our cohort into five bins corresponding to increasing PTV burden with a nearly equal number of individuals in each. Mean age for the bins was 57.7, 57.5, 57.5, 57.4, 57.4 years, respectively, with no difference in age distribution across the groups (Kolmogorov-Smirnov test on 2 samples p-value> 1%).

The Kaplan-Mayer survival curves for each group are shown in Figure 2, further highlighting elevated mortality of the subjects with higher PTV burden, with the most significant difference between cohorts #1 and #5 (log-rank test *p* = 7.1 × 10^−5^).

**Figure 2:**
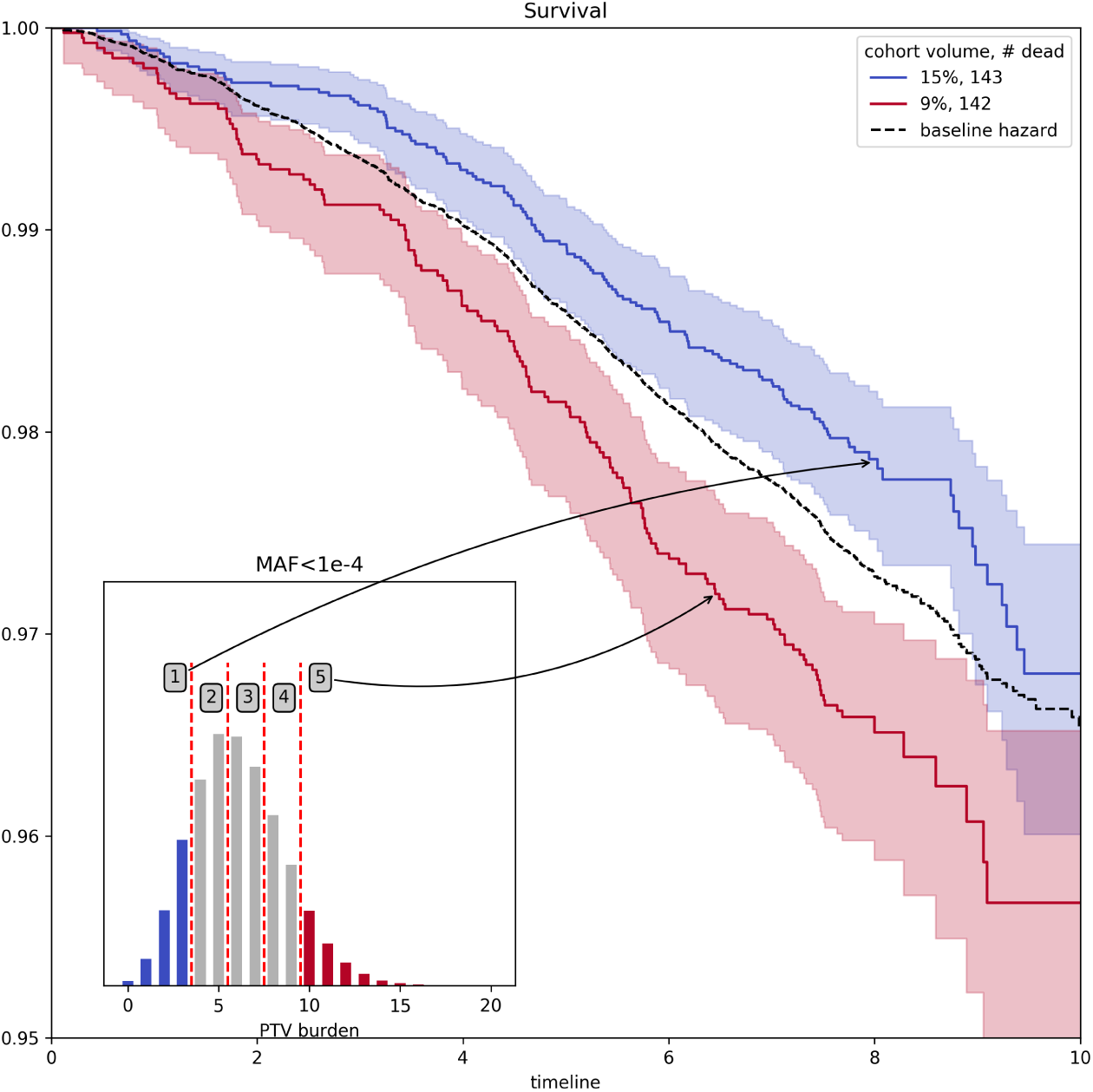
Survival curves for the cohorts from UKB stratified into five bins of increasing ultra-rare (*MAF* < 0.0001) PTV burden. Bins with lowest and highest PTV burden are labeled in blue and red, respectively, in the inset, and the corresponding survival curves are shown in the figure.

Having established the association of PTV burden with lifespan, we explored other major genetic variant types selected for incidence frequency and category: 3-prime and 5-prime UTR region variants, TF binding sites and structural interaction variants (Table S2). Of all tested PTV types, the most significant associations with lifespan and healthspan were observed for the ultra-rare (*MAF* < 0.0001) stop gain, splice donor and frameshift variants (Figure 3). However, only stop gains were associated with mother’s age at death and none of the categories were associated with father’s age at death. As a negative control, we also included the effects of neutral variants - synonymous gene variants, showing no significant effect on our longevity phenotypes.

**Figure 3:**
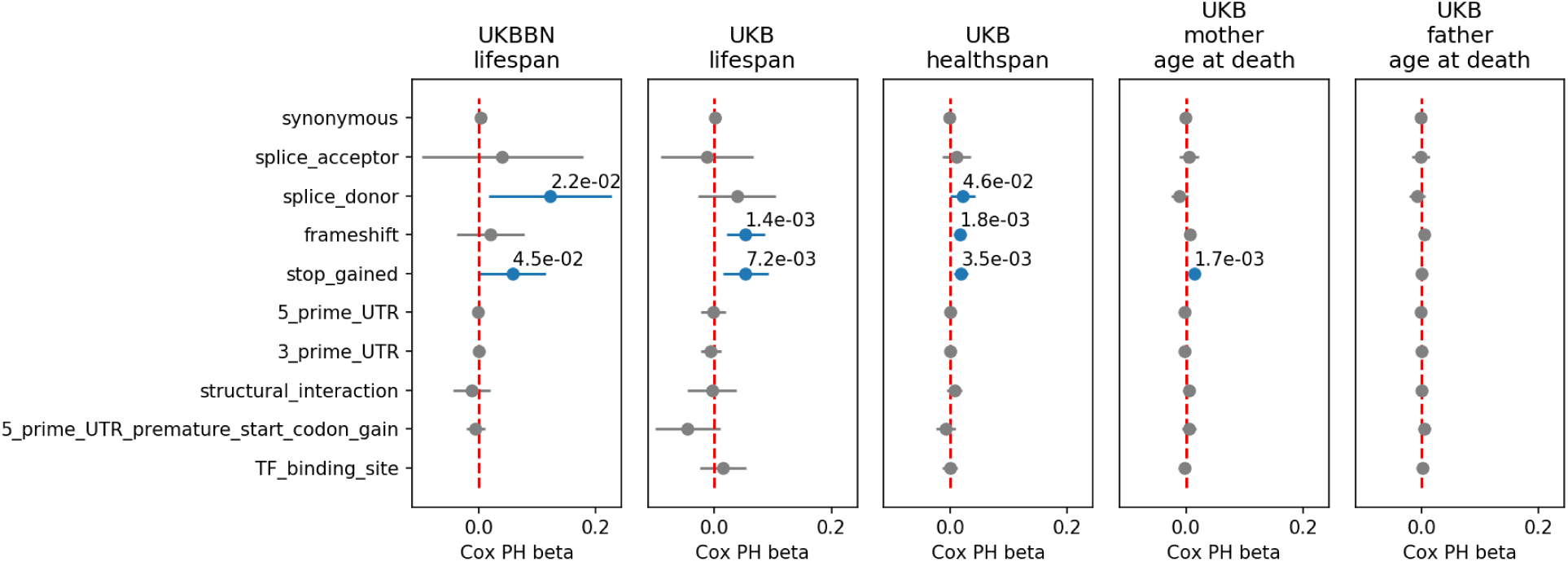
Association of ultra-rare (*MAF* < 0.0001) and putatively damaging gene variant burden and the UKB and UKBNN lifespan, UKB healthspan, and parental longevity (father’s and mother’s age at death).

### Gene constraint analysis

Although ultra-rare PTVs affect nearly 90% of sequenced genes in UKB dataset, there were genes intolerant to rare PTV (iPTV) that harbored none of the ultra-rare PTVs (*MAF* < 0.0001) in whole population. We compared these genes with those harbring at least one PTV (*n* = 16, 495) within the same 4 MAF bins tested for the association with lifespan. The unaffected (iPTV) group of genes on average was expressed in more tissues (Figure 4a) and had higher indispensability scores (a metric to measure gene essentiality introduced by^18^, Figure 4b). In line with this finding, genes that harbored more frequent PTVs on average were more tissue-specific and had lower indispensability scores, thus were less essential, in agreement with previously published results reported for ExAC cohort^9^. Moreover, ultra-rare stop gain variants were more likely to trigger NMD (Figure 4c) as predicted by snpEff based on the 50 bp rule^19^ and in line with what was previously demonstrated for rare variants in GTEx dataset^7^.

**Figure 4:**
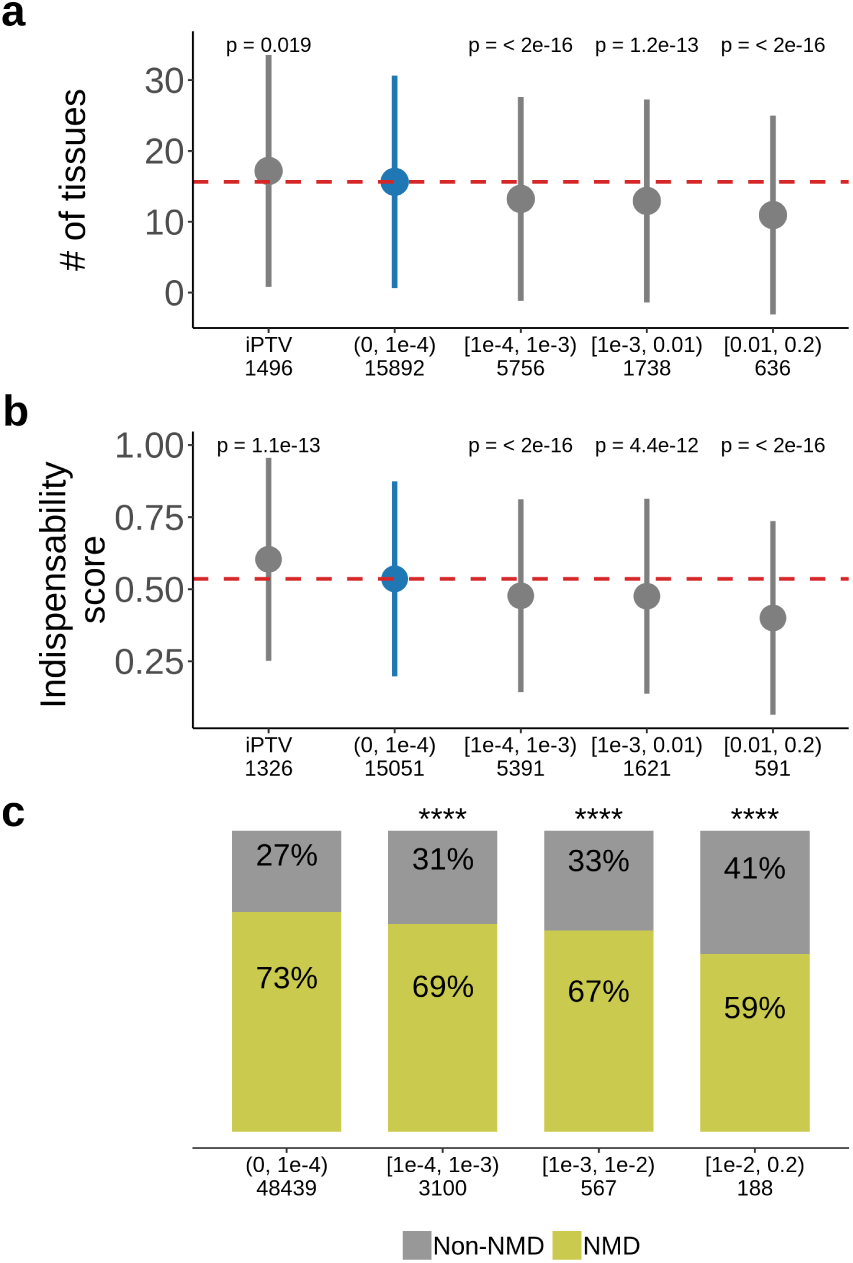
Characteristics of genes harboring PTVs of different MAF bins or lacking PTVs (iPTV) in UKB population. Numbers below each bin represent the number of genes harboring PTVs of the corresponding MAFs included in the analyses. PTV-intolerant (iPTV) and genes harboring rare PTVs (see (0, 10^−4^) bin) **a** are more broadly expressed and **b** have higher indispensability scores (a metric to measure gene essentiality introduced by ^18^). The results of comparisons are grouped in subsequent MAF bins and the numbers in the horizontal axis represent the number of genes included in the analysis. **c**, Rare stop gains are more likely to trigger NMD. Numbers in the horizontal axis correspond to the total number of stop gains included in the analysis. Each group was compared to the bin (0, 10^−4^), marked blue, where PTVs are significantly associated with lifespan. P-values in **a** and **b** calculated by Wilcoxon rank-sum test, p-value in **c** calculated by Fisher exact test, **** p-value < 0.0001

We further hypothesized that subjects with the same number of PTVs may have different lifespan due to difference in the damaging effect of their PTVs, wherein subjects dying earlier harbor more deleterious alleles that those dying later in life. To test this idea, we compared gene characteristics of subjects with the same PTV number (5 PTVs per exome, *n* = 171) but different lifespan (Figure 5a). Subjects dying younger harbored more damaging PTVs compared to the subjects with longer lifespan. PTVs in shorter-lived subjects resided in the genes characterized by higher genome-wide haploinsufficiency score (GHIS^20^) (Figure 5b). They also affected more broadly expressed genes based on RNA sequencing data from GTEx dataset (Figure 5c). Also, these genes were more intolerant to PTVs due to the lower observed/expected (oe, gnomAD v2.1) scores as in the gnomAD these genes harbored fewer PTVs than expected. Moreover, mean values for constraints of the genes disrupted by PTVs showed association across lifespan tested by Cox PH model. Percent of tissues expressing these genes, oe scores and loss-of-function occurrence were all significantly associated with lifespan within group of subjects with 5 PTVs (Figure 5a).

**Figure 5:**
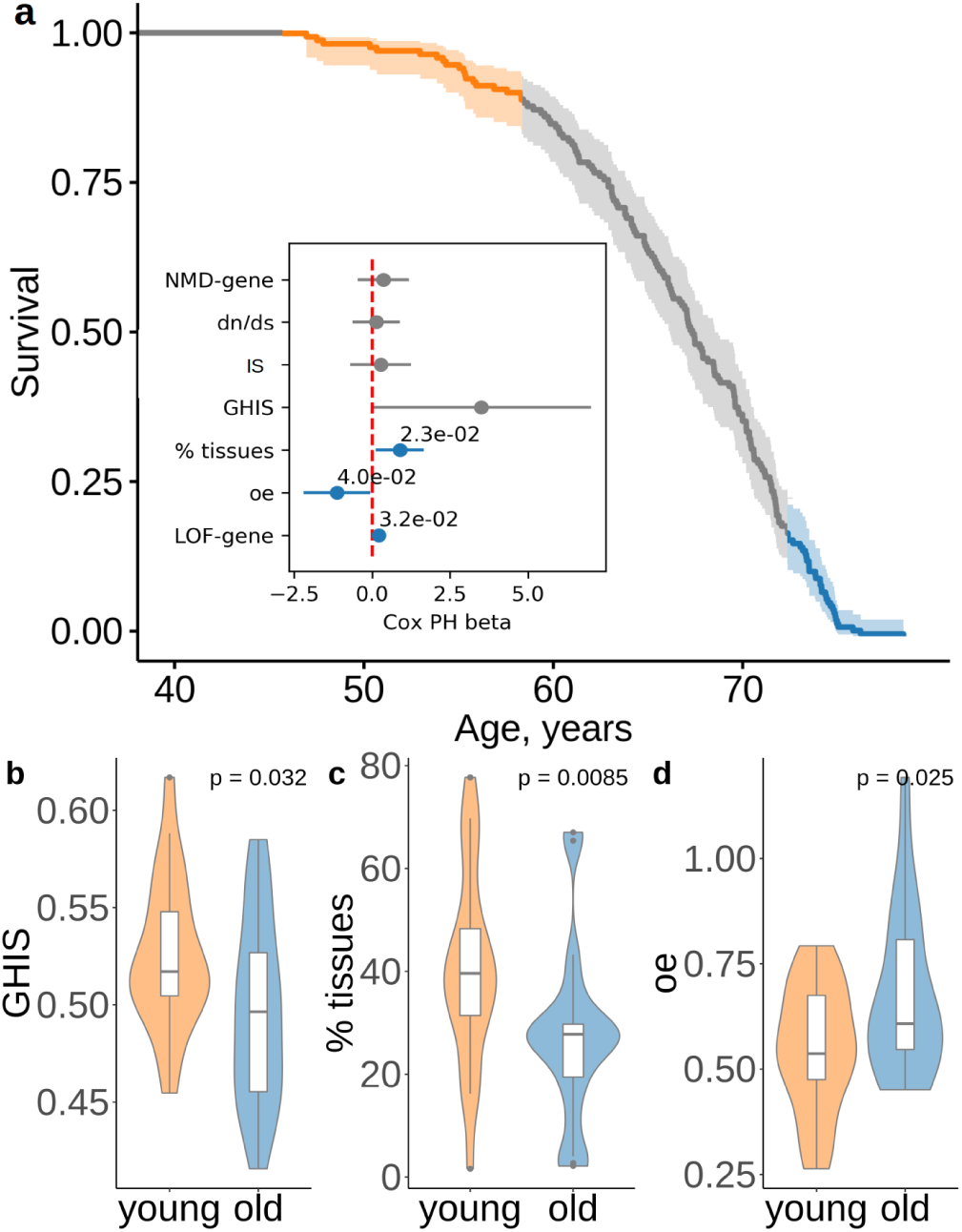
**a**, Survival of subjects with 5 ultra-rare PTVs per exome in UKB. The inset shows the association between lifespan and gene variant properties: evolutionary constraint quantified by *dN/dS* ratios (the ratio of substitution rates at non-synonymous and synonymous sites) in human-chimpanzee orthologs; indispensability score (IS) as in ^18^; genome-wide haploinsufficiency score (GHIS) as in ^20^; (relative) number of tissues expressing the gene; observed/expected (oe) score; prediction for variants being loss-of-function (LOF, see LOF-gene) and triggering NMD (see NMD-gene). Orange and blue areas in **a** designate survival windows for subjects dying earlier in life (young) and later in life (old), and this color scheme is the same as that in the plots **b**-**d**. Difference in **b**, GHIS scores, **c**, percent of tissues expressing gene affected by variants, and **d**, oe scores for individuals with same PTV number but differing in lifespan (i.e. dying younger (47.4 − 58.9 years) or older (73.8 − 78.5 years)). P-values in **b** and **d** are calculated by Student t-test, and p-value in **c** is calculated by Wilcoxon rank-sum test.

### Somatic mutations and mortality acceleration

In addition to the germline burden of extremely rare PTVs, somatic cells stochastically accumulate new genetic variants^21–23^ at a median mutation frequency, *R* ≈ 10^−8^ per base pair per year^24^. Thus, the negative effect on healthspan and lifespan due to germline burden should get gradually amplified with age in somatic cells. We quantitatively assessed whether the contribution of stochastically accumulated PTVs is strong enough to explain the exponential growth of mortality with age, a.k.a. the Gompertz law. In doing so, we extrapolated the Cox PH model for germline PTV burden by taking into account the effects of acquiring new PTVs with age in somatic cells. The somatic PTV burden increases linearly with age and can be estimated as *λLRt*, where *t* is age, the genome size is *L* = 3Gbp, and the fraction of the genome covered with the extremely rare PTVs (10 kbp) with *MAF* < 10^−4^ is *λ* = 0.33 · 10^−5^. Overall, the somatic PTV burden contributed to the mortality log-hazard a linear (in age) term *βλLRt*, where *β* = 0.046 per year is the Cox PH coefficient, whereas the Gompertz contribution would be proportional to Γ*t*. We estimate that the somatic PTV burden term *βλLR* ≈ 4.6 · 10^−6^ per year is negligible in comparison to the Gompertz exponent Γ ≈ 0.09 per year characterizing mortality and incidence of chronic disease acceleration with age^4^. The estimated effect of somatic PTV accumulation is minor in comparison to the effect of germline PTV burden, and can account only for a minute fraction of mortality and morbidity acceleration.

## Discussion

We report that both lifespan and healthspan are negatively impacted by the burden of ultra-rare PTVs. These mutations are harbored by most humans, and thus disease occurrence and age of death are directly influenced by the genotype that defines a person at conception. Our approach is radically different from the previous searches for common variants that contribute to longevity, e.g. searches for alleles enriched in centenarians. In this regard, we find that the extended lifespan may be explained by both certain common variants, as found by previous studies, and by lacking ultra-rare damaging PTVs that reduce lifespan.

These conclusions are based on the analysis of two independent datasets. Due to the limited follow-up, mortality in the UKB dataset reflects the progression rate of age-related chronic diseases in an individual; i.e. if a subject is deceased, he/she most probably had one or more age-related diseases at the time of enrollment. The association of ultra-rare PTV mutations with healthspan, however, reflects the effect of deleterious gene variants on the incidence of the first chronic disease and thus covers the accumulated effect of genotype on health and survival over a much longer time, effectively from birth up to the age of enrollment/death.

The association of the ultra-rare mutation burden and longevity in the UKB dataset, which has more subjects but narrower age distribution, is consistent with the result of the analysis of the UKBNN cohort, which provides postmortem genotypes of individuals deceased at ages of 16−105 years old. By the nature of its design, the UKBNN cohort may be enriched for individuals prone to diseases and death at any given age. On the other hand, UKB subjects exhibit lower mortality and hence are probably healthier than the general population^25^. It, therefore, appears that the association of the ultra-rare mutations and longevity is a general feature that applies to diverse populations.

The ultra-rare PTV burden was also associated with a reduced mother’s but not father’s age at death. Aside from being a useful longevity proxy in UKB, through shared genetic makeup, parental survival effectively offers an independent population comprised of UKB participants’ parents for the validation in our study. The small effect size may be due to the fact that the effects of parental genes are diluted in the following generation^16^.

The relation between frequency of mutations and their association with the longevity traits is not trivial. For example, if an effect of a mutation does not depend on mutation frequency, the significance of association between the mutation incidence and the corresponding trait should increase as the mutation frequency increases. Therefore, the inverse effect-to-frequency relation for the frequency of PTVs and their effect on lifespan and healthspan in our analysis may be a signature of purifying selection for longevity.

Ultra-rare PTVs occur across the genome and affect 89% of sequenced genes in UKB. Intriguingly, we observed a subset of 1, 496 genes that are free of ultra-rare PTVs in the whole UKB population. These genes are characterized by higher indispensability scores and expressed in more tissues. Overall, these iPTV genes are more essential than the genes harboring PTVs. Together with their absence in all 40K participants, this finding indicates strong purifying selection against PTVs in these genes. Their disruption could lead to either childhood or embryonic lethality, the time periods that are not covered by our datasets. It is apparent that the rare PTVs affecting lifespan found in our study are not too damaging to cause early life mortality.

As expected, there were fewer genes affected by more frequent PTVs (*MAF* > 10^−4^). These genes are less conserved (Figure S3), less essential (based on indispensability scores) and expressed in fewer tissues. Moreover, stop gains corresponding to more frequent MAF bins were less likely to trigger NMD. Overall, more frequent PTVs apparently affect expression less and disrupt less essential genes. This indicates that these PTVs should have lower effect on fitness, explaining the lack of the association between the burden of more frequent PTVs and lifespan.

Analysis of rare PTVs in subjects with the same PTV number but different lifespan revealed that individuals with shorter lifespan harbor more deleterious alleles. This indicates that variability in lifespan in those subjects can be attributed to the quality of ultra-rare PTVs as well as environmental factors. These alleles reside in more broadly-expressed genes, affecting 15 tissues on average, and in genes that are more intolerant to PTVs (according to gnomAD oe scores) compared to the longer living group. At the same time, these variants are more likely to be loss-of-function. Thus, both the degree of damage caused by each variant and the number of these variants are important.

The combined effect of ultra-rare deleterious mutations was comparable to the effect of frequent mutations, previously associated with longevity based on human GWAS. For example, *ϵ*4 allele in *APOE/TOMM40* locus conferred an estimated 1.24 years of life lost (women only), as inferred from a large parental survival study^16^. The PTV number difference of 2 − 3 variants at the standard deviation for *MAF* < 0.0001 observed in this study, corresponds to the same order of effect (1 − 2 years) of lifespan and healthspan variation, under the Cox PH model.

Taken together, the effects of common variants earlier implicated in longevity and the effects of ultra-rare variants reported here could help explain the apparent heritability of lifespan. Currently, this issue is not fully resolved. Twin studies^26, 27^ suggest that lifespan could be as much as 23 − 33% genetically determined. A more recent study^5^ puts up a challenge to that conclusion and points to a much lower level of genetic determinism. We therefore expect that future investigations of the effects of ultra-rare genetic variants may turn to be crucial for quantitative understanding of lifespan heritability.

Having established the mortality and morbidity risk association with PTV variants, we were able to factor in the characteristic rates of mutation accumulation in human genome over the lifespan. The dramatic discrepancy between the estimate for the effect of accumulation of somatic PTV burden under the Cox PH model and the empirical mortality and morbidity acceleration does not support the hypothesis that random somatic mutations significantly reduce health- or lifespan. Moreover, the calculation shows that the effect of random accumulation of somatic mutations is less profound than that of germline PTV burden. We thus found little evidence for a significant role of somatic mutations in aging^28, 29^. Somatic mutations may, however, play a role through high-order effects, such as clonal expansion and cell competition triggered by PTVs, and hence amplify the effects of other forms of damage^30^. These findings strengthen the case for complexity of aging, wherein aging is a systemic process resulting from the combined accumulation of age-related deleterious changes, none of which could cause aging on their own^31^. The advantage of mutations in aging studies, however, is that they can be quantified and their contribution estimated, which is something that is currently much more difficult to do for other forms of age-related damage.

## Methods

### UKB cohort

The first batch of UKB exome sequence group consists of 49, 960 individuals who passed QC procedures by UKB. Exome sequencing cohort is enriched with samples with a higher rate of imaging and enhanced measurements such as retinal optical coherence tomography test, visual acuity, hearing test, and other. This cohort is not biased on any health condition, disease or physical measurement results from the UKB population of almost 500 thousand individuals^32^. We selected a cohort of 41, 250 individuals who self-reported ‘White British’ and have very similar genetic ancestry based on a principal components analysis of the genotypes. Then, we made an effort to produce the maximal independent set of individuals based on computed kinship coefficients (two individuals were considered related if they share relatedness of third degree or closer) and selected 40, 368 individuals for the analysis.

### Exome data

Exome data consisted of 8, 959, 608 SNPs and short indels from human coding DNA. We selected 6, 208, 943 variants that are not monomorphic in our cohort of interest and have a missing rate less than 10% and *MAF* < 0.2. We annotated these genetic variants for functional consequence using SNPeff^33^ software and *GRCh38.86* genome reference. UKKBN dataset was additionally annotated with ANNOVAR^34^ to add ExAC MAFs. Then, we defined protein-truncating variants as having any of the major damaging consequence: stop codon gained, frameshift variant, slice donor or acceptor variant, this produces 152, 790 and 11, 393 SNPs and indels in UKB and UKBBN, respectively, that we used for further analysis.

### PTV burden calculation and Cox proportional hazards model

PTV burden was defined as the number of ultra-rare (*MAF* < 10^−4^) variants that disrupt open reading frame (stop gain, frameshift, splice donor/acceptor). PTV burden was tested for association with UKBBN lifespan using Cox PH model with sex and first 20 principal components as covariates in R^35^. Eigen vectors were obtained from variants with *MAF* > 10% pruned using 50 window size, step size of 5 and variance inflation factor threshold of 1.5 by Plink^36^.

For UKB data we included sex, all available 40 genetic principal components and assessment centers as covariates for Cox PH analysis on lifespan, healthspan and mother’s and father’s age at death. For all types of survival data except for healthspan we’ve also added age at assessment as covariate. Genetic principal components were calculated on genotypes data for 500, 000 UKB participants^2^.

### UKBBN PCA with 1000G

First and second chromosome for all 1000G super populations were clustered with the UKKBN dataset. For that, 1000G vcf files were lift over to hg19 using picard tools, combined with UKBBN vcf file by overlap variants using GATK tools. Variants with MAF deviating between datasets over 30% were excluded. Eigen vectors were obtained from variants with *MAF* > 10% pruned using 50 window size, step size of 5 and variance inflation factor threshold of 1.5 by Plink. PCA plots were visualized in Rstudio. We kept individuals that clustered with European population (Figure S2).

### Data filtering

PTVs in UKB were filtered using internal MAFs. Since UKBBN cohort is much smaller to get desired resolution we used ExAC MAFs for non-finish European population (`ExAC_NFE`). We excluded ultra-rare variants absent in ExAC dataset (`ExAC_ALL` = 0) from UKBBN analysis to reduce number of sequencing and variant calling errors. Analysis in both datasets was restricted to autosomal chromosomes to avoid sex bias. We restricted UKKBN cohort to natural causes of death (i.e. excluding car accidents, poisoning and suicides) and excluded deaths with no abnormalities detected.

### Data sources

UKBBN vcf files were downloaded from EGA repository (EGAS00001001599, https://www.ebi.ac.uk/ega/studies/EGAS00001001599). Transcripts per kilobase million (TPM) counts for 53 human tissues were downloaded from GTEx Portal, release v7. Gene expression values within brain regions, two heart and two skin samples were averaged for subsequent analysis, and primary cell cultures were excluded yielding 37 tissues. Transcripts considered to be expressed in the tissue if *TPM* > 10. Oe ratios were downloaded from gnomAD repository (gnomad.v2.1.1.lof_metrics.by_gene.txt.bgz). GHIS values were obtained from^20^ and indispensability scores were downloaded from^18^. *dS* and *dN* values for chimpanzee-human orthologs were downloaded from Ensembl Biomart. NMD and LoF predictions were obtained from snpEff annotation (‘NMD.gene’, ‘LoF.gene’)^33^. All UK Biobank data are available upon application.

## Acknowledgements

This research has been conducted using the UK Biobank Resource under Application Number 21988. The study was funded by NIH (to V.N.G.) and by Gero LLC. The provision of UKBBN data used in this study was supported by funding from the UK Medical Research Council and BDR (Brains for Dementia Research). The authors thank Prof. Dmitry Ivankov for a helpful discussion.

## Authors’ contributions

V.N.G., P.O.F. designed the research; A.V.S., A.A.Z., A.E.T. analyzed the data; A.V.S., A.A.Z., A.E.T., P.O.F., V.N.G. wrote the manuscript.

## supplementary material

**Table S1:** Incidence of first disease (end of healthspan) statistics.

|  | Number of events | % |
| --- | --- | --- |
| cancer | 6239 | 53.3 |
| diabetes | 2009 | 17.2 |
| MI | 1862 | 15.9 |
| COPD | 619 | 5.3 |
| stroke | 527 | 4.5 |
| dementia | 211 | 1.8 |
| death | 126 | 1.1 |
| CHF | 114 | 1.0 |

**Table S2:** Variant annotations for 8, 959, 608 SNPs from FE dataset. Variant types selected for analysis marked in italics and PTV burden components marked in bold. Some variants may have multiple effects.

| Variant Effect | Number of variants |
| --- | --- |
| intron_variant | 3643472 |
| missense_variant | 2281322 |
| synonymous_variant | 1159078 |
| splice_region_variant | 333226 |
| downstream_gene_variant | 329399 |
| upstream_gene_variant | 303346 |
| *3_prime_UTR_variant* | 303346 |
| *5_prime_UTR_variant* | 192159 |
| ***frameshift_variant*** | 96359 |
| intragenic_variant | 85619 |
| sequence_feature | 79868 |
| ***stop_gained*** | 68054 |
| *structural_interaction_variant* | 57365 |
| *TF_binding_site_variant* | 45909 |
| *5_prime_UTR_premature_start_codon_gain_variant* | 34381 |
| ***splice_donor_variant*** | 22476 |
| disruptive_inframe_deletion | 21392 |
| ***splice_acceptor_variant*** | 18591 |
| conservative_inframe_deletion | 12612 |
| disruptive_inframe_insertion | 11080 |
| intergenic_region | 11012 |
| conservative_inframe_insertion | 8665 |
| start_lost | 5807 |
| stop_lost | 2442 |
| protein_protein_contact | 1590 |
| stop_retained_variant | 1077 |
| initiator_codon_variant | 609 |
| TFBS_ablation | 180 |
| bidirectional_gene_fusion | 17 |
| gene_fusion | 7 |
| exon_loss_variant | 5 |
| 3_prime_UTR_truncation | 3 |
| non_canonical_start_codon | 2 |

**Figure S1:**
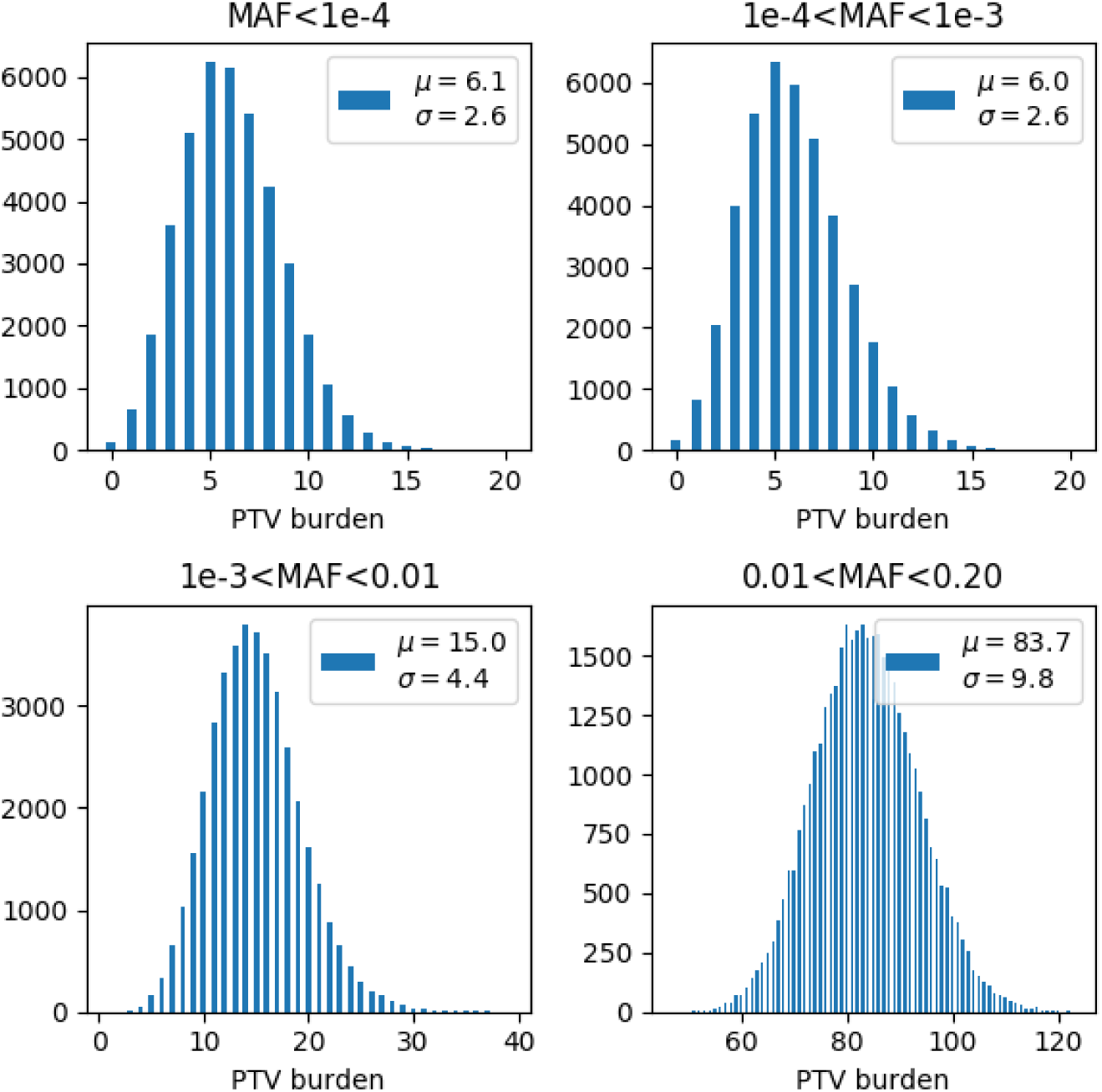
Histograms of PTV burden distributions depending on the PTV frequency.

**Figure S2:**
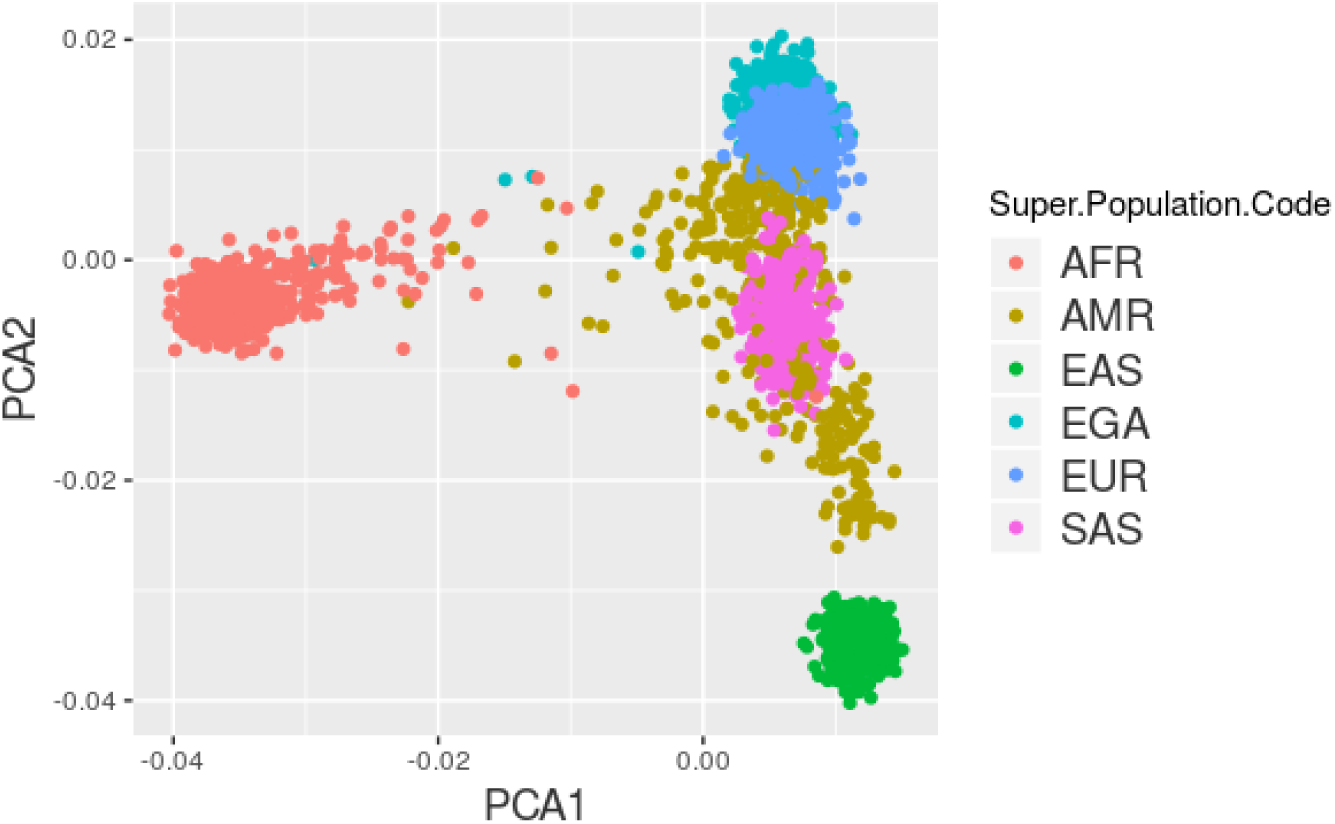
Population structure of the UKBBN dataset. PCA plot representing combined UKKBN (here designated as EGA) and 1000 Genome datasets restricted to 1st and 2nd chromosomes. AFR — Africans, AMR — Americans, EAS — East Asians, EUR — Europeans, SAS — South Asians.

**Figure S3:**
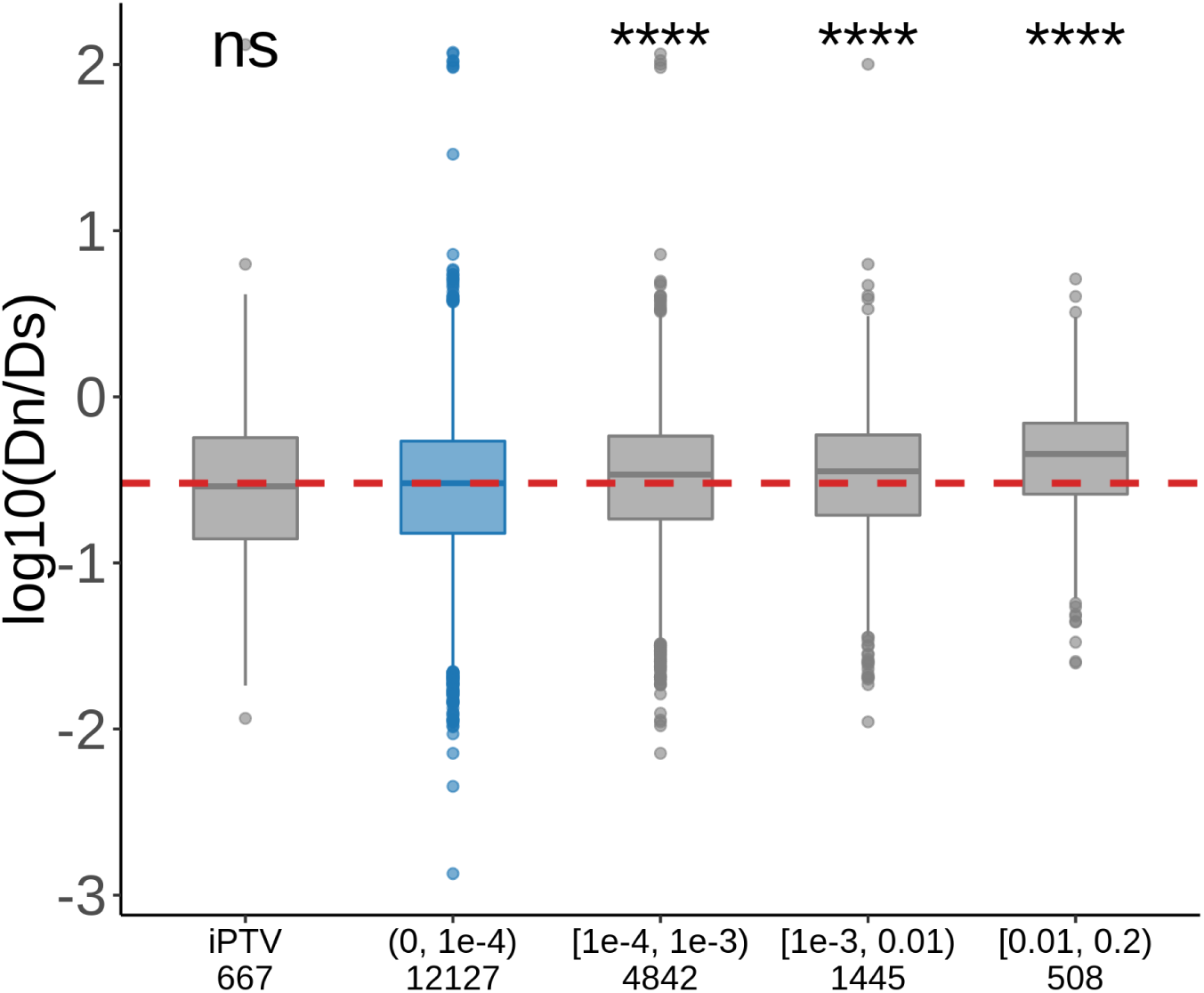
Human-chimpanzee *dN/dS* ratios for genes harboring PTVs belonging to different MAF bins or lacking PTVs (iPTV) in UKB population. Numbers below each bin represents number of genes harboring PTVs of corresponding MAFs included in the analyses.

